# Immediate and carryover reproductive costs of infection in female but not male house finches

**DOI:** 10.64898/2026.08.06.741781

**Authors:** K.M. Talbott, A.E. Fleming-Davies, F.E. Tillman, C.M.V. Nuñez, J.N. Weil, A.A. Perez-Umphrey, D.M. Hawley, J.S. Adelman

## Abstract

Wildlife diseases cause well-documented and often dramatic reductions in host survival. However, the impact of infectious diseases on host reproduction remains understudied, especially with respect to effects of prior and/or current pathogen exposure on reproductive development. Here we experimentally tested how prior and/or current infection with a common bacterial pathogen, *Mycoplasma gallisepticum* (‘MG’), alters reproductive development for female versus male house finches (*Haemorhous mexicanus*). Finches were inoculated with either MG or sterile media while in wintering condition and subsequently received one of these treatments while in breeding condition. In females, MG exposure had both immediate and carry-over effects on reproduction: controls had higher odds of laying eggs compared to females inoculated with MG in spring only, higher odds than females inoculated in both winter and spring, and higher odds than females given MG in the winter only. Conversely, breeding-condition males inoculated with MG in spring had higher testosterone levels than males receiving only control inoculations, and there were no carryover effects of winter MG inoculation or inoculations during both seasons on testosterone. Sex bias in the reproductive impacts of infectious diseases may have important knock-on effects on the epidemiology and population-regulating effects of pathogens, thereby warranting further study.

## Introduction

Infectious diseases threaten the viability of wild vertebrate populations worldwide [1]. Outbreaks frequently cause mass mortality events, yet the full impact of infectious diseases on host populations may not be clear from mortality alone. Infections acquired during the breeding season can also directly reduce host reproduction (e.g., [2–5]), while those acquired outside of the breeding season may have carryover effects (e.g., [6–8]) on future reproduction. Compared to survival costs, the reproductive costs of wildlife diseases have received less attention, yet our ability to make predictions about transmission dynamics and host population recovery [9] relies on accurately estimating the impact of emerging diseases on both axes of fitness.

Infectious diseases are known to decrease wildlife reproduction. For example, North American bat maternity colonies had fewer reproductive females when white nose syndrome was present [10]. Similarly, highly pathogenic avian influenza outbreaks were associated with nest failure in several bird species [11,12]. While later-stage reproductive impacts such as fetal loss [2–5,13] or offspring abandonment [2,14,15] are most apparent in the wild, pathogens might also reduce host reproduction prenatally due to trade-offs in investment between reproduction and self-maintenance [16]. For example, shifts in endocrine dynamics [17–19], delayed initiation of breeding [20–22], and decreased gamete production and/or quality [23–26] may carry important fitness consequences but are difficult to measure in sexes in situ. Controlled experiments are needed to investigate these early-stage reproductive costs, helping us better understand the full reproductive impact of wildlife diseases.

Such impacts may also vary by seasonal timing of infection. While reproductive impacts of infections acquired during the breeding season may be most obvious, some pathogens are transmitted year-round. This creates the potential for carryover effects of infections acquired outside of the breeding season on reproductive development for one or both sexes; however, few studies have explicitly examined these carryover effects in wildlife [6,7,10]. Reproductive impacts of infections acquired during non-breeding periods might differ from those acquired during the breeding season, due to shifts in physiological priorities. For example, some seasonal breeders preemptively limit self-maintenance costs during breeding by reducing standing immune defenses or immune responsiveness [27–29], with the type and extent of these fluctuations varying by sex [28,30–33]. Thus, experiments investigating the impact of infectious diseases on reproductive output, and trade-offs between reproductive and immune function, should control for both host sex and physiological state (i.e., breeding or non-breeding) during pathogen exposure.

We addressed how pathogen exposure during either non-breeding (‘wintering’), breeding (‘spring’), or both states impacts reproductive development in wild-caught house finches (*Haemorhous mexicanus*) experimentally inoculated with the bacterial pathogen, *Mycoplasma gallisepticum* (‘MG’). MG spilled over from domestic poultry in eastern North America during the 1990s, causing population declines in wild finches [34,35]. Now endemic, the pathogen continues to limit survival [36] by impeding anti-predator and foraging behavior [37–40]. Recovered finches harbor partial but incomplete immune protection and therefore reinfections occur readily [41–43]. While MG circulates year-round, prevalence peaks during autumn, following the influx of naive juveniles into the population, and again during mid to late winter [44]. The latter peak, occurring right before the breeding season, may be most relevant for knock-on effects on reproduction. Thus, to understand the reproductive costs of MG exposure, we must consider the interaction between physiological state and MG exposure history in finch reproductive development.

To address these objectives, we experimentally manipulated daylength in wild-caught finches to shift physiological state [45,46] and inoculated birds with either MG or a carrier control on two occasions. We investigated the effect of treatment on reproduction, using egg laying and circulating testosterone levels as proxies of reproductive development in females and males, respectively. Additionally, we examined the relationship between infection severity and reproductive development and asked whether this relationship varied by host sex.

## Methods

### Experimental design

We varied both winter and spring exposure to MG in a fully factorial design, resulting in four treatment combinations: Control Twice, Winter MG Only, MG Twice, or Spring MG Only (Table 1). Finches were first held on a winter (i.e., non-breeding) photoperiod and inoculated with either MG (18 males, 12 females) or a carrier control (20 males, 10 females); see supplementary methods for further inoculation details. Five weeks later, finches were shifted to a spring (i.e., breeding; see below) photoperiod (modified from [47]) for the remainder of the experiment. After four weeks on a spring photoperiod, finches were inoculated again either with MG (12 females, 20 males) or a carrier control (10 females, 18 males). See Figure S1 for timeline details and supplementary methods for information on finch capture, care, and sexing.

**Table 1.** House finches were assigned to one of four treatment regimens that each included one inoculation with *Mycoplasma gallisepticum* (‘MG’) or a carrier control while finches were in winter (i.e., non-breeding) condition and another inoculation while finches were in spring (i.e., breeding) condition. Treatments are ranked from lowest to highest infection severity during spring inoculation, with severity represented by treatment-level means of maximum individual MG loads, eye score, and anti-MG IgY levels.

| Treatment (n) | Mean Spring MG load ± SE | Mean Spring eye score ± SE | Mean Spring IgY ± SE |
| --- | --- | --- | --- |
| Control twice (n = 14) | 0 ± 0 | 0 ± 0 | < 0.01 ± < 0.01 |
| Winter MG only (n = 14) | 5 ± 2 | 0 ± 0 | 0.03 ± 0.01 |
| MG twice (n = 16) | 1663 ± 1656 | 0.28 ± 0.25 | 0.04 ± 0.01 |
| Spring MG only (n = 16) | 23645 ± 3056 | 2.06 ± 0.36 | 0.10 ± 0.02 |

### Data and sample collection

On days -1, 0, 3, 7, 14, 21, 28, 42, and 49 following winter inoculation, and days -1, 3, 7, 14, 21, and 28 following spring inoculation, we collected eye pathology scores (‘eye scores’). Eye scores were assessed for each eye on an ordinal scale and reflect the degree of conjunctival swelling, from no visible swelling (score of 0) to the eye swollen shut (score of 3); scores for both eyes are then summed [48]. We also collected blood for quantifying anti-MG IgY [49] in all birds and circulating testosterone in males; see Figure S1. On the last day of sampling, we inspected female cages for eggs and female abdomens for evidence of a brood patch, a featherless patch skin used for egg incubation [50]. See supplementary methods for further details on sample collection and analysis.

### Statistical analyses

We used ordered ranks to score female reproductive development: we assigned the highest rank to females that produced eggs and showed evidence of a brood patch, followed by females with only a brood patch, and finally by females with no brood patch and no evidence of egg laying. Male reproductive development was quantified by changes in testosterone levels, given by subtracting the highest observed circulating testosterone level (in ng/mL) before spring inoculation from the highest observed level following spring inoculation. Individuals with positive values showed an increase in circulating testosterone following spring inoculation.

To investigate the effect of treatment on female reproductive rank, we used ordinal regressions (*polr* function from the ‘MASS’ package in R [51]). Beta estimates were exponentiated for interpretation as odds ratios. For males, we used Kruskal-Wallis Rank Sum tests to ask whether testosterone change varied by treatment and used a Dunn’s test to compare differences between treatments. Model fit and assumption testing details are in the electronic supplementary material. We determined significance as p < 0.05.

We used a multivariate framework to estimate relationships between treatment, reproduction, and highly correlated inoculation response metrics. To reduce the dimensionality of inoculation response data (winter maximum MG load, spring maximum MG load, winter maximum eye score, spring maximum eye score, winter maximum IgY level, and spring maximum IgY level), we conducted a non-metric multidimensional scaling analysis (NMDS, stress value < 0.06) and computed the Bray-Curtis dissimilarity index to reduce our dataset to two dimensions (*metaMDS* function and vegan package in R [52]). We used a permutational multivariate analysis of variance (PERMANOVA) to investigate the effect of treatment on our resulting distance matrix (*adonis2* function) and conducted post-hoc tests to determine pairwise differences between treatments using the *pairwise.adonis* function. Finally, we asked whether NMDS 1 and/or NMDS2 predicted variation in reproductive metrics for each sex by comparing ordinal regression models (females) or linear models (males). We used AICc for model comparison, which corrects for small sample sizes [53], and selected models with the lowest AICc as the best performing models. All analyses were conducted in R [54].

## Results

### MG inoculation reduced reproductive development in females but not males

Experimental MG exposure, especially in breeding-condition females, reduced the probability of females reaching the highest reproductive rank (developing a brood patch and laying eggs; ordinal regression LR = 19.16, p < 0.01, McFadden’s R^2^ = 0.42). Females given two control treatments were 9.88 x 10^16^ times more likely to reach the highest reproductive rank compared to females given MG only in spring (odds ratio = 1.01 x 10^-17^, 1.01 x 10^-17^ – 1.01 x 10^-17^ CI; Figure 1A and Table 2). Similarly, females given two control treatments were 81 times more likely to lay eggs than females given two MG inoculations (odds ratio = 1.23 x 10^-2^, 4.88 x 10^-4^ – 0.31) and 29 times more likely to lay eggs than females receiving MG only during winter (odds ratio = 3.42 x 10^-2^, 1.42 x 10^-3^ – 0.83). Females given two MG inoculations also had lower MG loads in spring compared to females inoculated only in spring (Table 1), likely due to the partial immune protection provided by the winter exposure [41,55].

**Figure 1.**
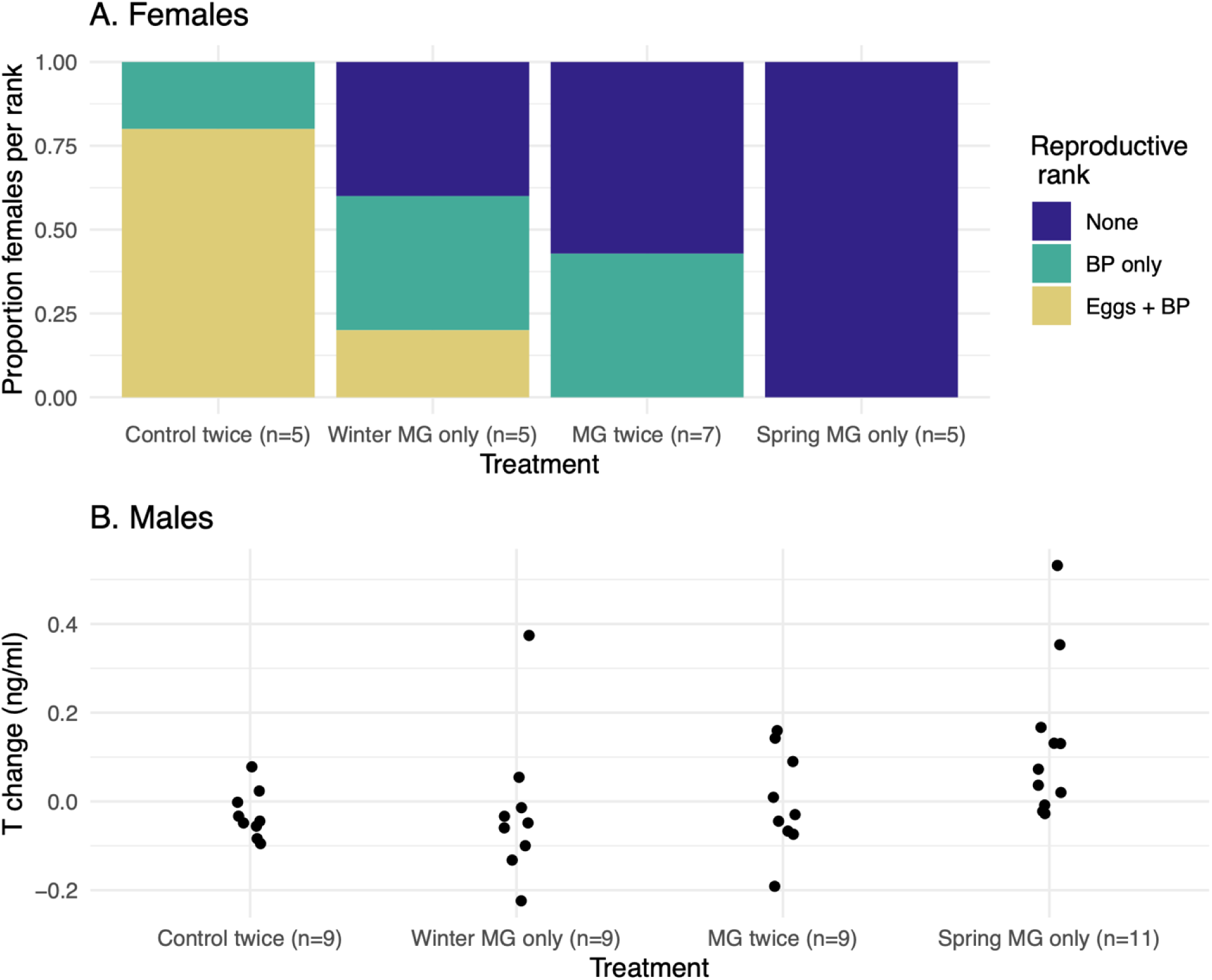
In house finches, inoculation with *Mycoplasma gallisepticum* (‘MG’) reduces female reproductive metrics but increases male testosterone production. Each bird was first inoculated with either MG or a carrier control while in winter (i.e., non-breeding) condition and again while finches were in spring (i.e., breeding) condition; treatments are ordered from lowest to highest spring infection severity (Table 1). In females (A), reproductive rank reflects reproductive development, with the highest rank assigned to females that developed a brood patch and laid eggs (yellow), followed by females that developed a brood patch (‘BP’) only (green), and the lowest rank was given to females meeting neither criterion (dark blue). In males (B), the change in circulating testosterone (‘T’) following spring inoculation is shown on the y-axis; positive values reflect males whose circulating testosterone increased, while negative values reflect males whose circulating testosterone decreased. Points represent data from individual male finches and are jittered along the x-axis for easier viewing.

**Table 2.**
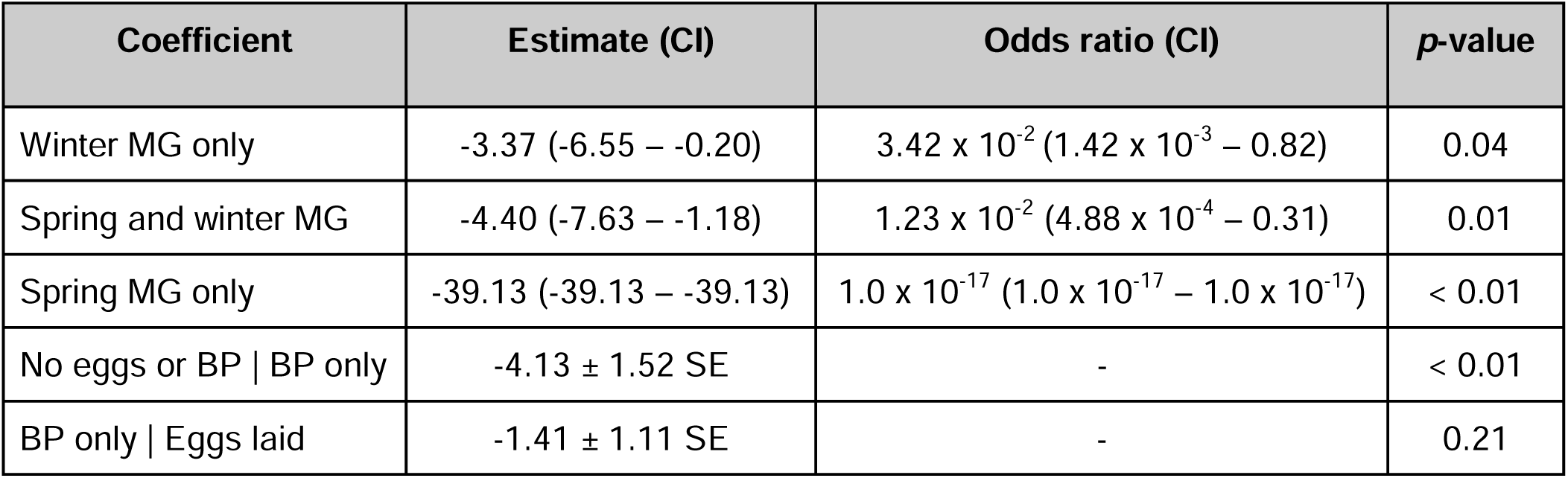
Coefficient table for an ordinal regression predicting female reproductive rank from treatment (n = 22). Females that developed a brood patch (‘BP’) and laid eggs were given the highest reproductive rank (‘Eggs laid’), followed by those that developed a brood patch but laid no eggs (‘BP only’), followed by those that met neither criterion (‘No eggs or BP’). Finches were inoculated with either *Mycoplasma gallisepticum* (‘MG’) or a carrier control on two occasions: while in winter (i.e., non-breeding) condition and while in spring (i.e., breeding) condition. Treatment regimens are ordered by their impact on spring infection severity from lowest to highest (see Table 1).

Male reproductive development, measured by testosterone change following spring inoculation, varied by treatment (Kruskal Wallis X^2^ = 9.80, df = 3, p = 0.02) in the opposite direction of females, with the largest testosterone increases in males with spring-only MG exposure (Figure 1B). Males treated with MG only in spring showed larger testosterone increases than males receiving two control treatments (Z = 2.54, adjusted p = 0.03) and males inoculated with MG only in winter (Z = 2.76, adjusted p = 0.04; Figure 1B), but there were no significant differences between other treatment pairs (p > 0.10).

### Inoculation responses varied by timing of MG exposure

The NMDS analysis of multiple infection metrics per individual (Figure 2) yielded two axes: axis 1 (i.e., NMDS1) is principally associated with severity of infection, with higher overall severity on the left, while axis 2 (i.e., NMDS2) is principally associated with timing of infection, with winter infections clustering toward the top and spring infections toward the bottom. Treatments clustered in different regions along these axes (PERMANOVA F = 57.81, df = 3, R^2^ = 0.76, p = < 0.01), with control animals clustered high along axis 1, those whose most severe responses occurred during winter (i.e., those inoculated in winter only or in both winter and spring) low along axis 1 and high on axis 2, and those whose most severe responses occurred during spring (i.e., those inoculated only in spring) clustered low along axis 1 and low along axis 2. Post-hoc analysis confirmed pairwise differences in inoculation responses (i.e., NMDS scores) between treatments, with the exception that finches inoculated with MG twice and those inoculated only in winter did not differ significantly (Table S2).

**Figure 2.**
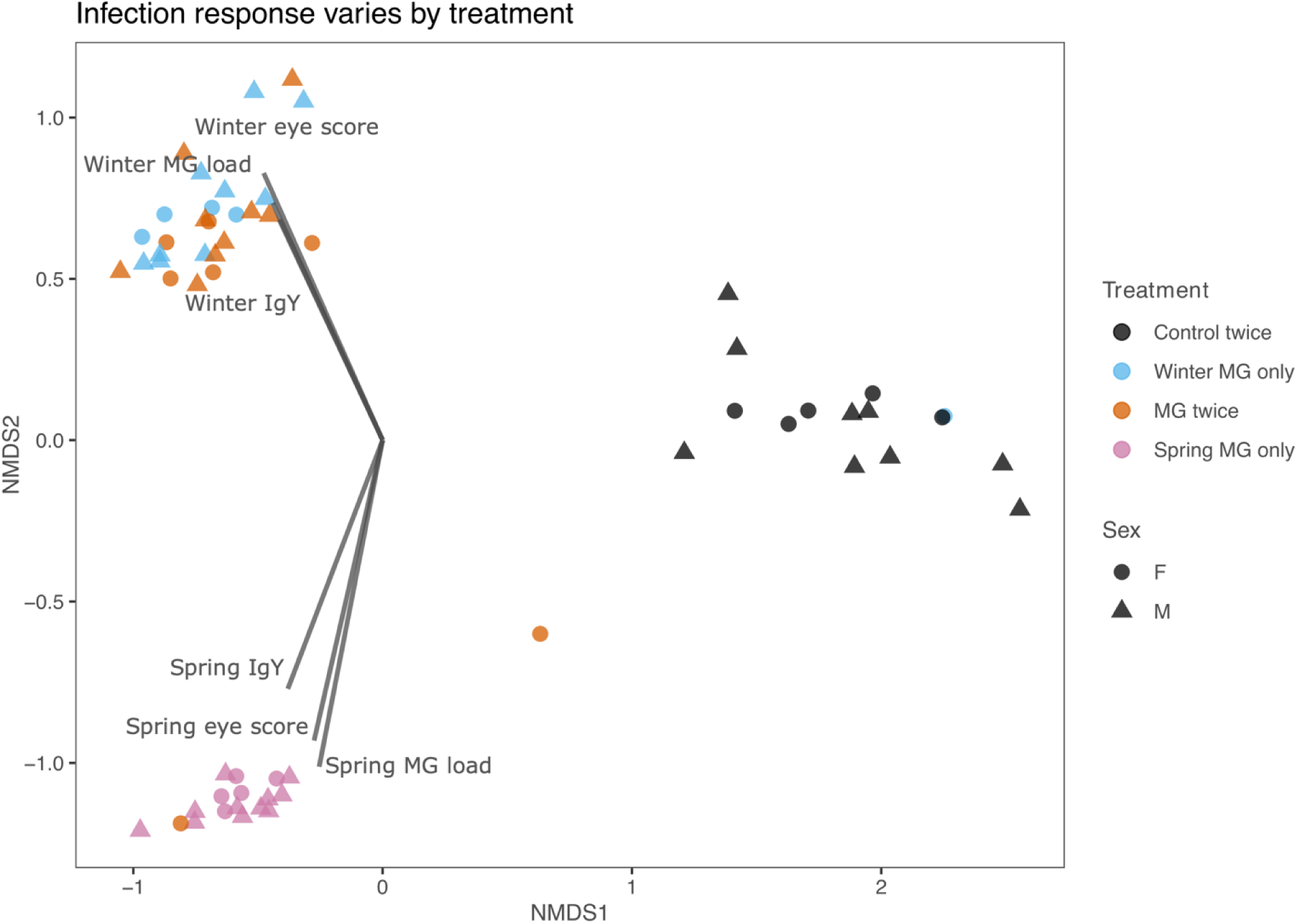
Ordination plot of two non-metric dimensions reflecting responses of house finches to different *Mycoplasma gallisepticum* (‘MG’) inoculation treatments. Six metrics of inoculation response (vectors labeled on ordination plot) were reduced to two dimensions using non-metric multidimensional scaling (NMDS). Finches were inoculated with either *Mycoplasma gallisepticum* (‘MG’) or a carrier control on two occasions: while in winter (i.e., non-breeding) condition and again while finches were in spring (i.e., breeding) condition, resulting in four treatment regimens (see Table 1). Points represent NMDS scores for individual finches. See Table S1 in the electronic supplementary material for loadings.

### Individual variation in disease response predicted reproductive development in females and males

Females with more severe overall inoculation responses (i.e., lower NMDS1 values), especially those whose most severe responses occurring during spring (i.e., lower NMDS2 values), had lower reproductive ranks, indicating a reduced probability of developing a brood patch and laying eggs; see Figures 3A, 3C, S4, and Table 3A. In contrast, males whose most severe inoculation responses occurred during spring (i.e., lower NMDS2 scores) showed relatively larger increases in testosterone. NMDS2 score alone was a better predictor of male testosterone change than models incorporating NMDS1 alone or both NMDS1 and NMDS2 (Figures 3B, 3D, S4B, and Table 3B).

**Figure 3.**
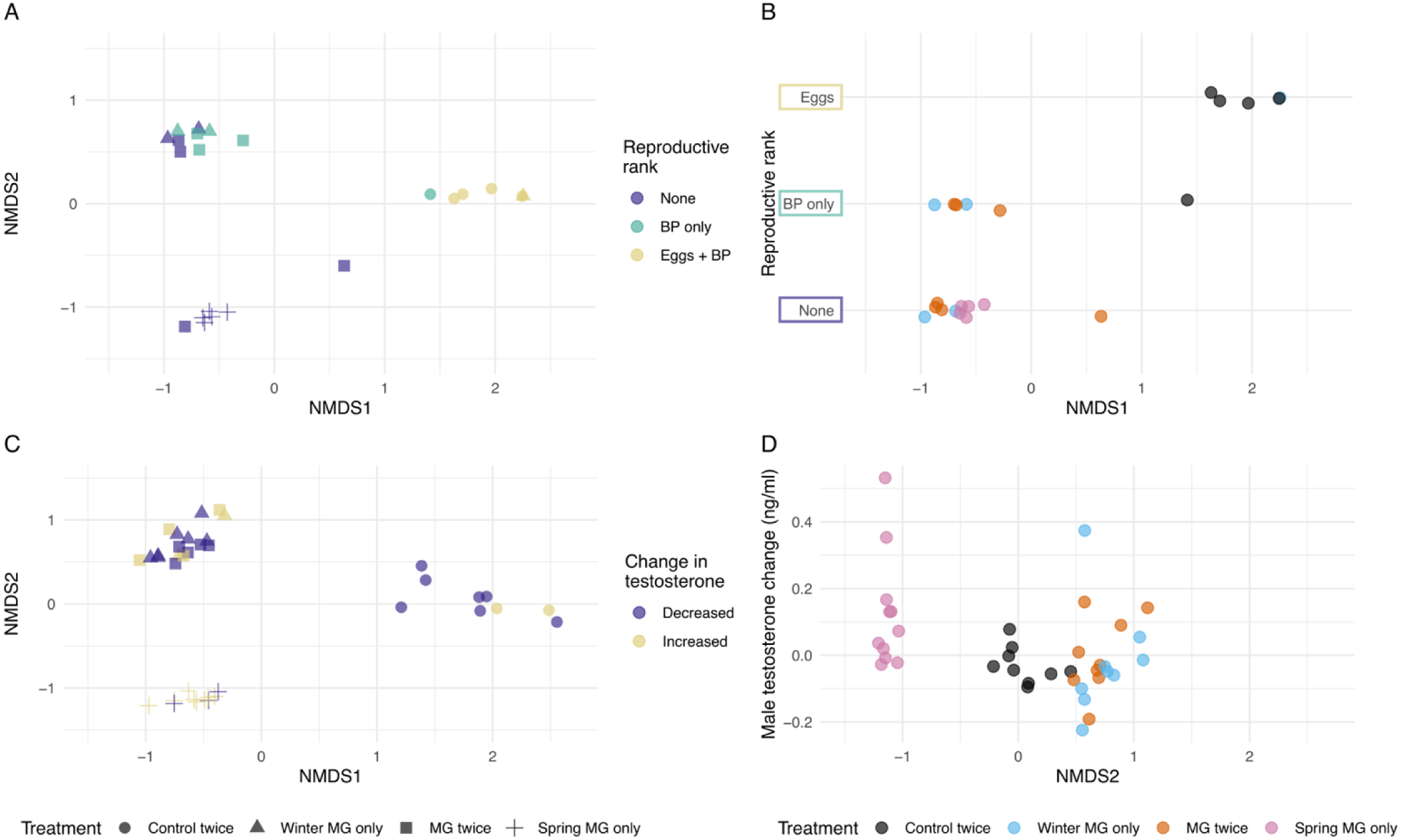
House finch reproductive metrics vary by *Mycoplasma gallisepticum* (‘MG’) treatment and inoculation response. Each bird was first inoculated with either MG or a carrier control while in winter (i.e., non-breeding) condition and another inoculation while finches were in spring (i.e., breeding) condition. Inoculation responses (maximum MG load, eye scores, and antibody levels) were collapsed to two axes by NMDS. Females (panels A and B) that developed a brood patch and laid eggs (“Eggs + BP”) were given the highest reproductive rank, followed by females only developing a brood patch (“BP only”) and females that met neither criterion (“None”); females given at least one MG inoculation, reflected by lower NMDS1 scores, were less likely to reach the highest reproductive rank than females receiving only control inoculations. Males (panels C and D) with the highest spring infection severity, reflected by more negative NMDS2 scores, showed higher increases in testosterone than males with lower spring infection severity; an increase in testosterone indicates an difference > 0 ng/ml in maximum post-spring inoculation testosterone levels minus maximum pre-spring inoculation levels, while a decrease in testosterone indicates a difference < 0 ng/ml. Points represent data from individual finches; see Figure S4 for additional plots of NMDS scores and reproductive metrics.

**Table 3.**
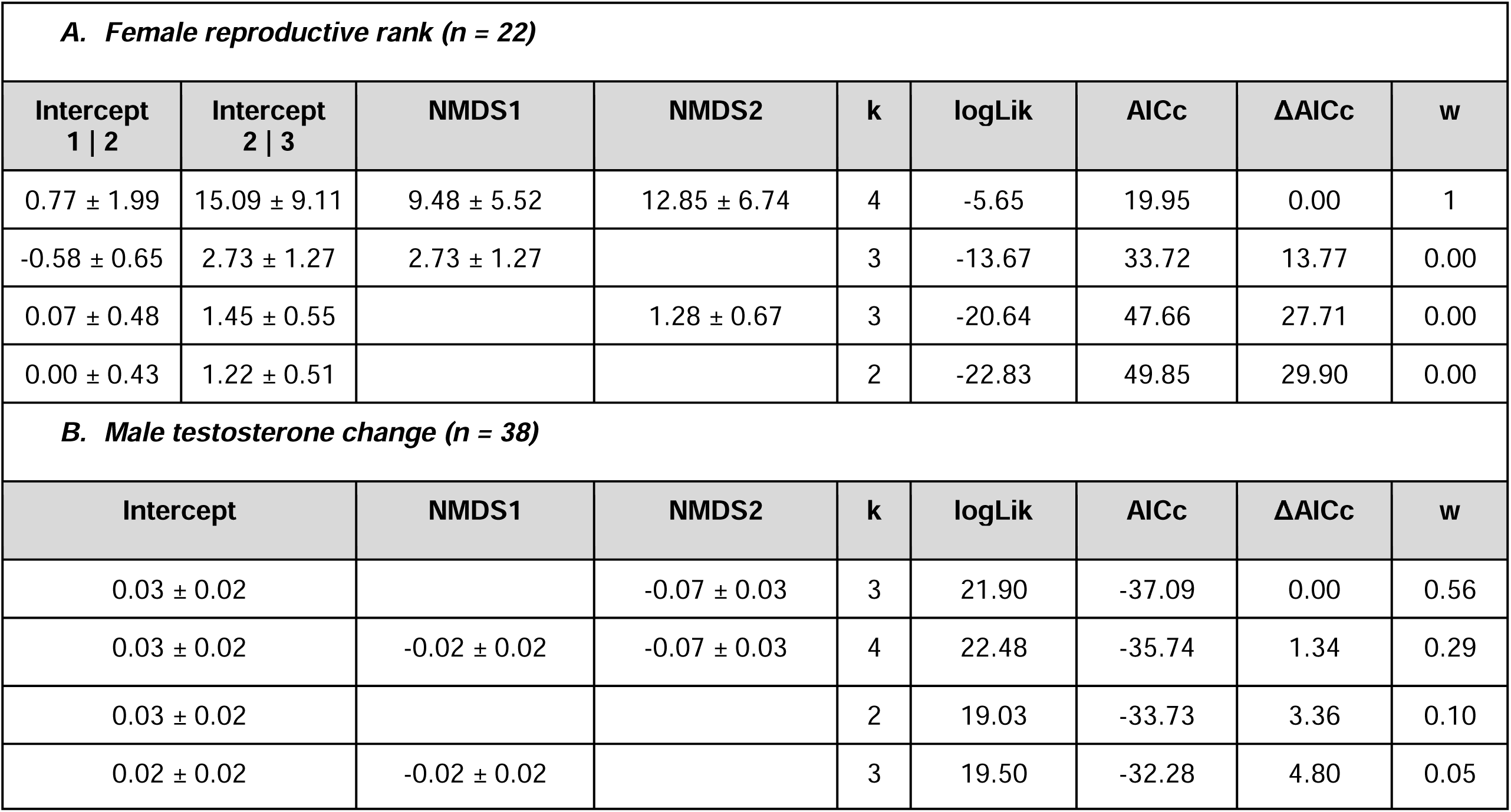
Models assessing predictors of reproductive development in house finches inoculated twice with *Mycoplasma gallisepticum* bacteria or carrier controls. Female reproductive development was quantified by a ranked variable with the highest rank assigned to females that developed a brood patch and laid eggs, followed by females only developing a brood patch, and then females meeting neither criterion. Male testosterone change (B) was quantified by the change in circulating testosterone following spring inoculation. Predictors include NMDS scores reducing the dimensionality of inoculation response data to two dimensions (NMDS1 and NMDS2). Beta estimates are given ± standard error.

## Discussion

Here we found evidence for both immediate and carryover costs of experimental pathogen exposure on female reproductive development (egg laying and brood-patch development) in wild-caught house finches. In contrast, males inoculated under breeding condition showed a larger increase in testosterone than winter-inoculated or uninoculated males. Although egg laying and testosterone production are not equivalent metrics of reproductive development, our results nonetheless suggest that the seasonal timing of pathogen exposure has important and sex-specific implications for reproductive fitness.

MG inoculation had the strongest negative reproductive impact on females inoculated only in spring, as demonstrated by treatment-level differences in reproductive rank, as well as the strong relationship between NMDS2 (a metric of infection severity by season) and reproductive rank. Because immune function and female reproduction are both metabolically costly [56–59], our results suggest that immune function was prioritized over reproduction in breeding-condition females. In addition, winter-only MG inoculation reduced the probability of egg laying, indicating that females experienced carryover effects of winter MG exposure on spring reproduction. Furthermore, females inoculated twice with MG also showed a reduced probability of egg laying; because most (n = 5 out of 7; see supplementary methods) were not successfully infected during spring, likely due to the acquired protection from winter inoculation, winter carryover effects likely reduced egg laying in this group.

Similar carryover effects occur in other systems, including reduced probability of pregnancy in female caribou (*Rangifer tarandus*) with heavy warble fly (*Hypoderma tarandi*) burdens during winter [7] and female North American bats in caves where white-nose syndrome was detected [10]. However, to our knowledge, our study is the first experimental demonstration of seasonal carryover effects of pathogen infection on host reproduction, with the exception of work in parasites that cause chronic infections (e.g., [20,60–62]). Given the documented carryover effects of other non-breeding season stressors (e.g., winter toxin exposure [63,64], habitat quality [65,66], and climate [67,68]) on wildlife reproductive success, pathogen infections acquired during the non-breeding season may represent a broader key influence on wildlife reproduction.

It is unclear whether non-laying females in our study were skipping reproduction or delaying reproductive development. Because brood patch development generally occurs shortly before egg laying [69], it is perhaps likely that non-layers with brood patches would have laid eggs eventually if the experiment had been continued. In either case, females exposed to MG for the first time during (or immediately prior to) their first breeding season may experience substantial reproductive costs, as delayed onset of breeding is generally associated with reduced reproductive success in birds [70–72]. Additional work is needed to fully quantify the reproductive costs of MG and determine the mechanisms by which infection might halt female reproduction, which likely include pleiotropic effects of endocrine or immune signaling [59].

Although male and female finches did not vary in infection severity, males showed distinct reproductive responses to MG treatment from females, with no evidence that MG inoculation reduced male reproductive development, as quantified by testosterone change. Sperm sampling would be a more direct measure of reproductive development to confirm this result. However, experimental inoculation with *Plasmodium* had no effect on sperm production in dark-eyed juncos (*Junco hyemalis*), a songbird with similar life-history patterns [73], suggesting it is unlikely that MG inoculation reduces house finch sperm production. Rather, the increased testosterone observed in spring-inoculated males could reflect a stronger investment in the current breeding season (i.e., terminal investment), stimulated by cellular damage caused by MG inoculation [74,75]). An alternative explanation is immunoredistribution, a hypothesized phenomenon in which testosterone, either directly or indirectly, diverts leukocytes to tissues where they are needed [76]. In this case, increased testosterone production could reflect a concerted immune response to address infection. Regardless of the underlying mechanism, our work suggests that males pay lower short-term reproductive costs of MG exposure compared to females. This is an important addition to the literature on impacts of pathogens on host reproduction, as reproductive indices are rarely assessed in both sexes separately.

Our results also have implications for transmission in wild populations. The influx of MG-naive juvenile finches at the end of the breeding season is thought to help drive seasonal epidemics [44,77], as is seen in other host-pathogen systems [78–81]. In the MG system, and likely others, naive individuals will generally have higher susceptibility to infection, higher loads, and more severe pathology than animals with immune protection from prior exposure, thereby increasing the role of juveniles in transmission [41,82,83]. The delayed onset of breeding in females with current or prior pathogen exposure could extend the breeding season, prolonging the influx of naïve juveniles, and potentially influence transmission. For males, elevated testosterone during the breeding season is associated with behaviors such as territoriality and courtship [84,85]; if pathogen exposure increases male testosterone instead of decreasing it, as shown in our study, infected males in the wild likely show typical reproductive behaviors that could increase transmission through physical contact. In the finch-MG system, this would reinforce earlier studies suggesting that adult male finches could disproportionately drive MG transmission [86,87]. Our study underscores the need for further work investigating sex differences in physiological responses to infectious diseases, which have the potential to impact both pathogen transmission and host population viability in wild vertebrates.

## Supporting information

Supplemental Analyses

## Acknowledgments

This project was supported by the National Institute of General Medical Sciences of the National Institutes of Health under Award Number R01GM144972 (to DMH, JSA, AEF, SJG, LMC, and KEL). We thank Anna Perez-Umphrey, Jesse Garrett-Larsen, Sara Teemer, Marissa Langager, Riley Meyers, Kayla Elahi, Cassie Sturgill, and Dean Hoff for assistance in finch capture and care at Virginia Tech. Thank you to Joshua Beer (University of Memphis) for assistance with animal handling and the University of Memphis Animal Care Facilities Staff for animal care. We thank Pierre Deviche for consulting on house finch testosterone EIA methods and Kate Boersma for consulting on statistical analysis.

## Data accessibility statement

All data and code are available at https://github.com/talbottkm/sex-differences-MG-finch-reproduction.

