## Supplemental Figures and Tables for "Immediate and carryover reproductive costs of infection in female but not male house finches"

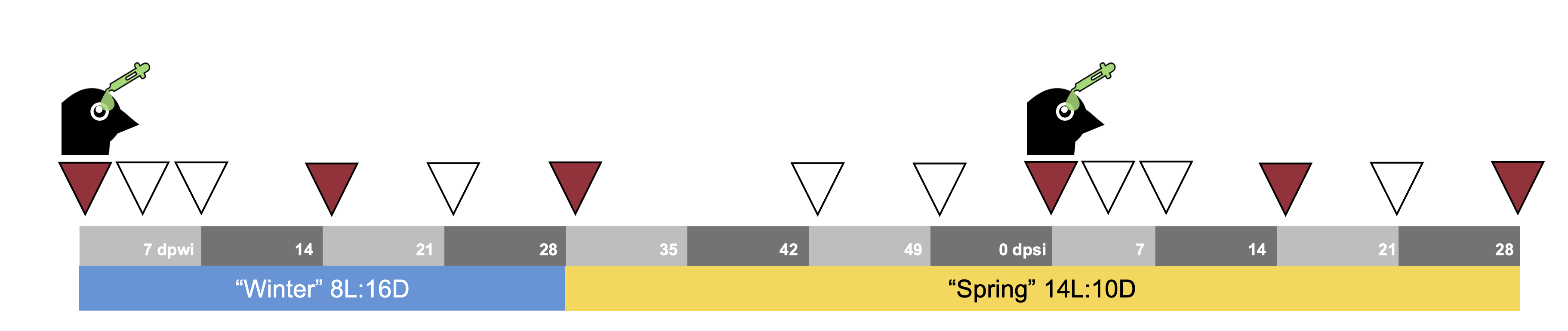

**Figure S1.** Experimental timeline. Birds with green droplets indicate dates that house finches were inoculated either with *Mycoplasma gallisepticum* or a carrier control. Finches were held on a winter photoperiod (8L:16D, hrs light:dark) for 8 wks prior to winter inoculation to induce a winter (i.e., nonbreeding) physiological state. After a further four weeks on a winter photoperiod, the photoperiod was then increased to stimulate a spring (i.e., breeding) physiological state. Gray bars indicate distinct weeks and each number gives the number of days post-inoculation that mark the end of each week (dpwi = days post winter inoculation, dpsi = days post spring inoculation). All triangles indicate dates of sampling eye pathology, MG load, and finch mass. On red triangle dates, blood was also sampled for antibody quantification in all finches and circulating testosterone levels in males (testosterone quantified only on 0, 14, and 28 dpsi). On the final sampling date (28 dpsi), females were checked for evidence of egg laying and brood patches.

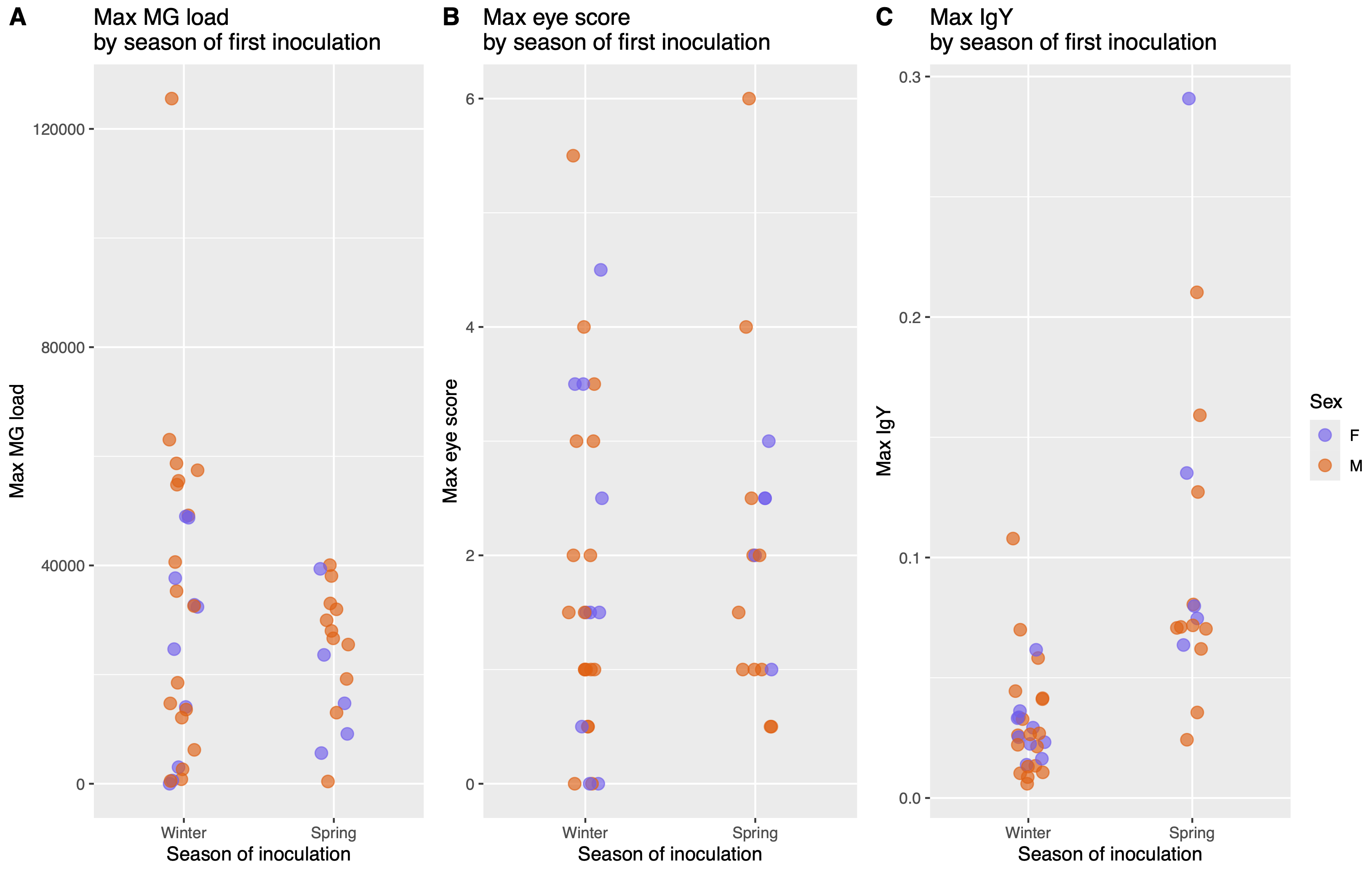

**Figure S2**. In house finches inoculated with *Mycoplasma gallisepticum* (‘MG’) for the first time, neither A) maximum load nor B) pathology (maximum eye score) varies by host sex or season of inoculation. However, C) maximum anti-MG IgY levels are higher in finches inoculated for the first time in spring (i.e., breeding) condition, compared to those inoculated for the first time in winter (i.e., non-breeding) condition. Points represent maximum values observed for individual finches that were successfully infected (n = 44); points are jittered along the x-axis for easier viewing. Female finches are shown in purple (n = 15) and males are shown in orange (n = 29).

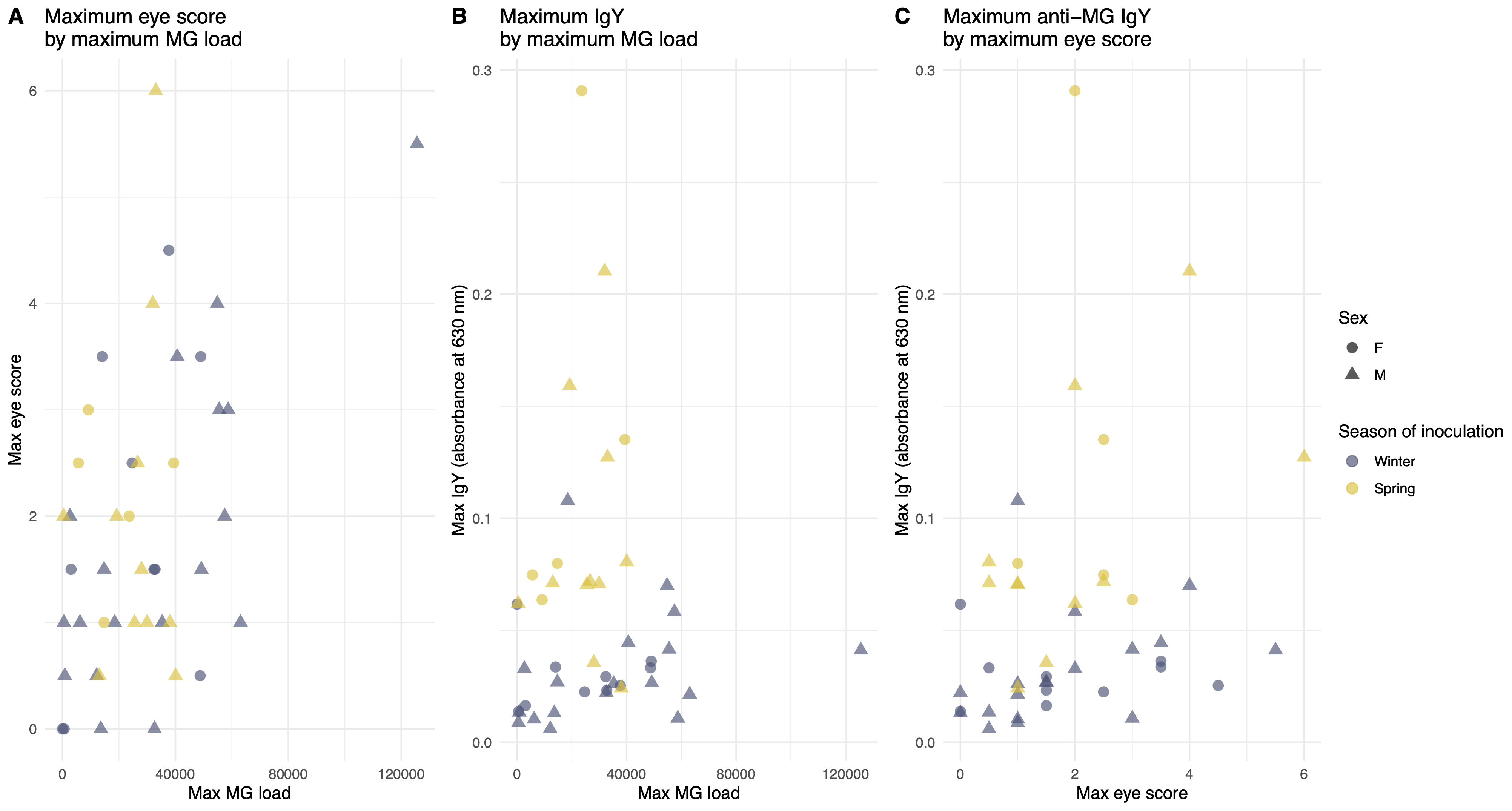

**Figure S3.** Shown are different metrics of inoculation response (eye pathology score, bacterial load, and antibody production) in house finches successfully inoculated for the first time with *Mycoplasma gallisepticum* (‘MG’). Points represent data from individual finches that were successfully infected with MG (n = 44), with females shown in round points (n = 15) and males shown in triangular points (n = 29). Finches inoculated for the first time while in winter (i.e., non-breeding) condition shown in dark blue, and those inoculated for the first time in spring (i.e., breeding) condition in yellow. See supplementary analyses for statistical relationships between inoculation responses.

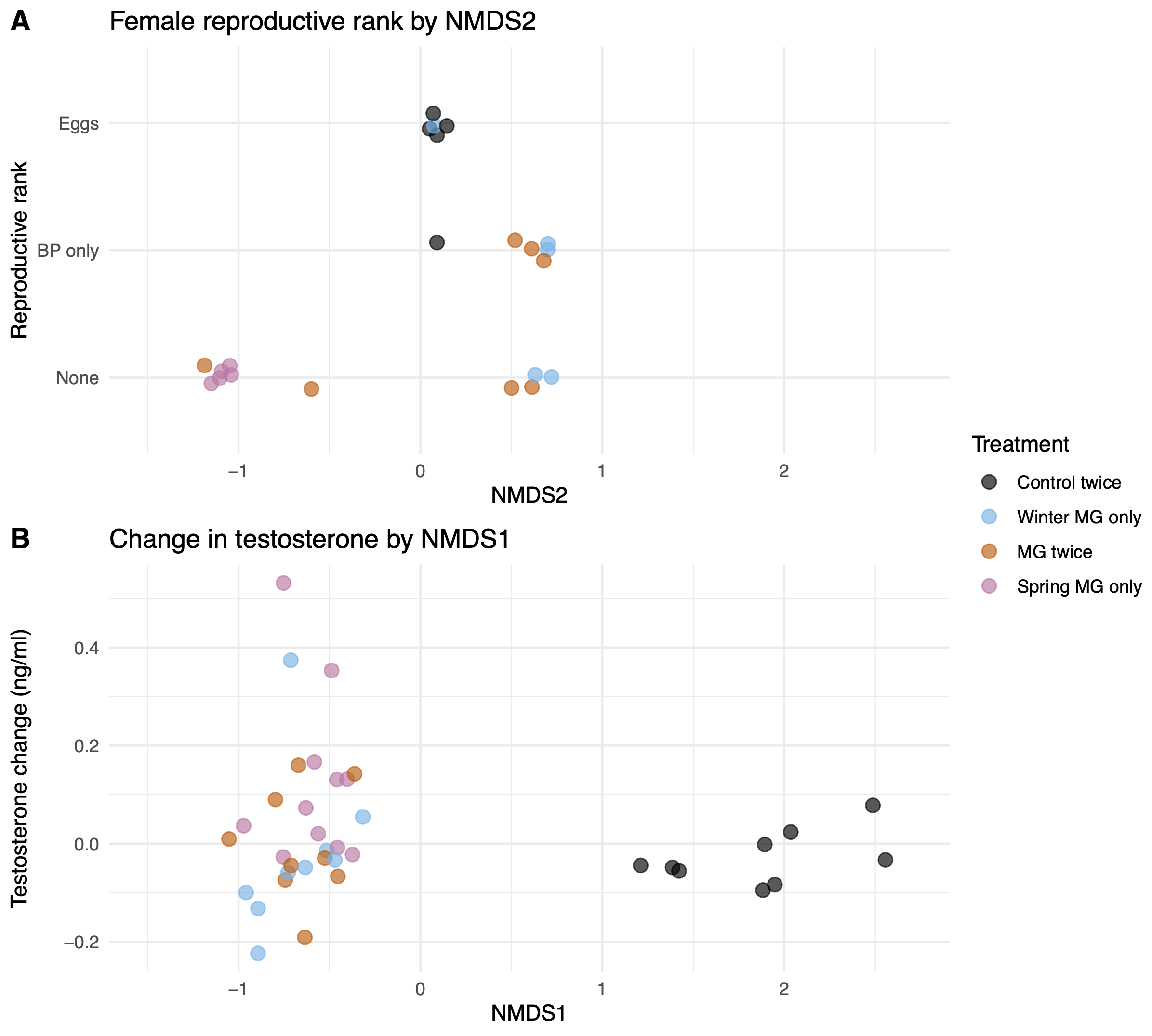

**Figure S4**. House finch reproductive metrics vary by *Mycoplasma gallisepticum* (‘MG’) treatment and inoculation response (n = 38 males, 22 females). Each bird was first inoculated with either MG or a carrier control while in winter (i.e., non-breeding) condition and again while finches were in spring (i.e., breeding) condition**.** Infection response data were reduced to two dimensions by using NMDS. NMDS1 scores reflect whether a bird received at least one MG inoculation (MG-inoculated finches have lower scores, while those given only control inoculations have higher scores), while NMDS2 reflects the season when a bird experienced the most severe infection (lower scores generally indicate severe infection during spring, while higher scores indicate severe infection during winter). A) Shown is the relationship between NMDS2 and female reproductive rank; the highest rank assigned to females that developed a brood patch and laid eggs, followed by females that developed a brood patch (‘BP’) only, and the lowest rank was given to females showing no signs of reproductive development. B) Shown is the relationship between NMDS1 and the change in circulating testosterone following spring inoculation; positive values reflect males whose circulating testosterone increased, while lower values reflect males whose circulating testosterone decreased. Points represent data from individual finches and female points are jittered along the y-axis for easier viewing.

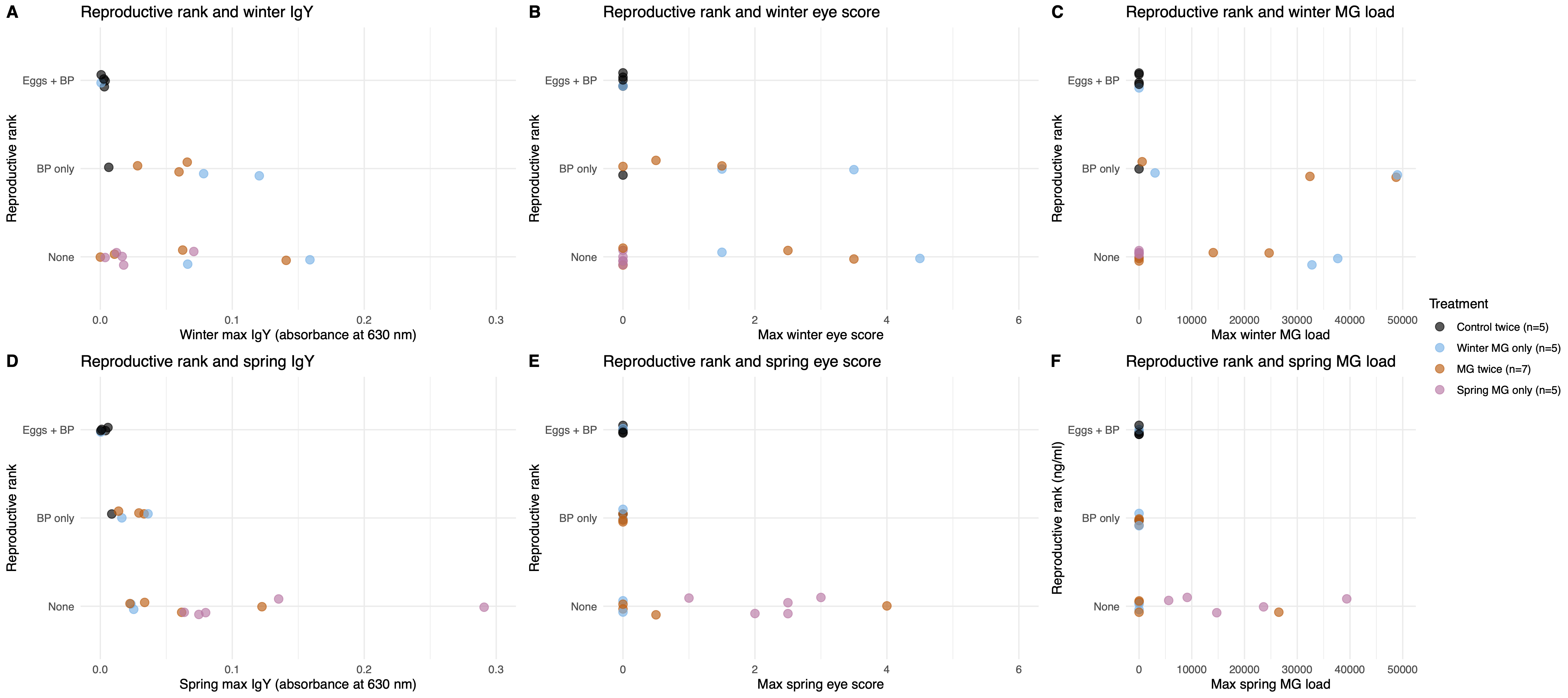

**Figure S5.** Relationships between female house finch (n = 22) reproductive rank and response to *Mycoplasma gallisepticum* (‘MG’) inoculation, shown by treatment regimen and inoculation response metric. Each bird was first inoculated with either MG or a carrier control while in winter (i.e., non-breeding) condition and then received a second inoculation while in spring (i.e., breeding) condition. The highest reproductive rank was assigned to females that developed a brood patch and laid eggs, followed by females that developed a brood patch (‘BP’) only, and the lowest rank was given to females showing no signs of reproductive development. Shown are relationships between reproductive rank and different seasonal metrics of inoculation response. Points represent data from individual finches and are jittered along the y-axis for easier viewing; points are colored by treatment, with finches given two control doses in black, those given MG only in winter in blue, those given two MG doses in orange, and those given MG only during spring in pink.

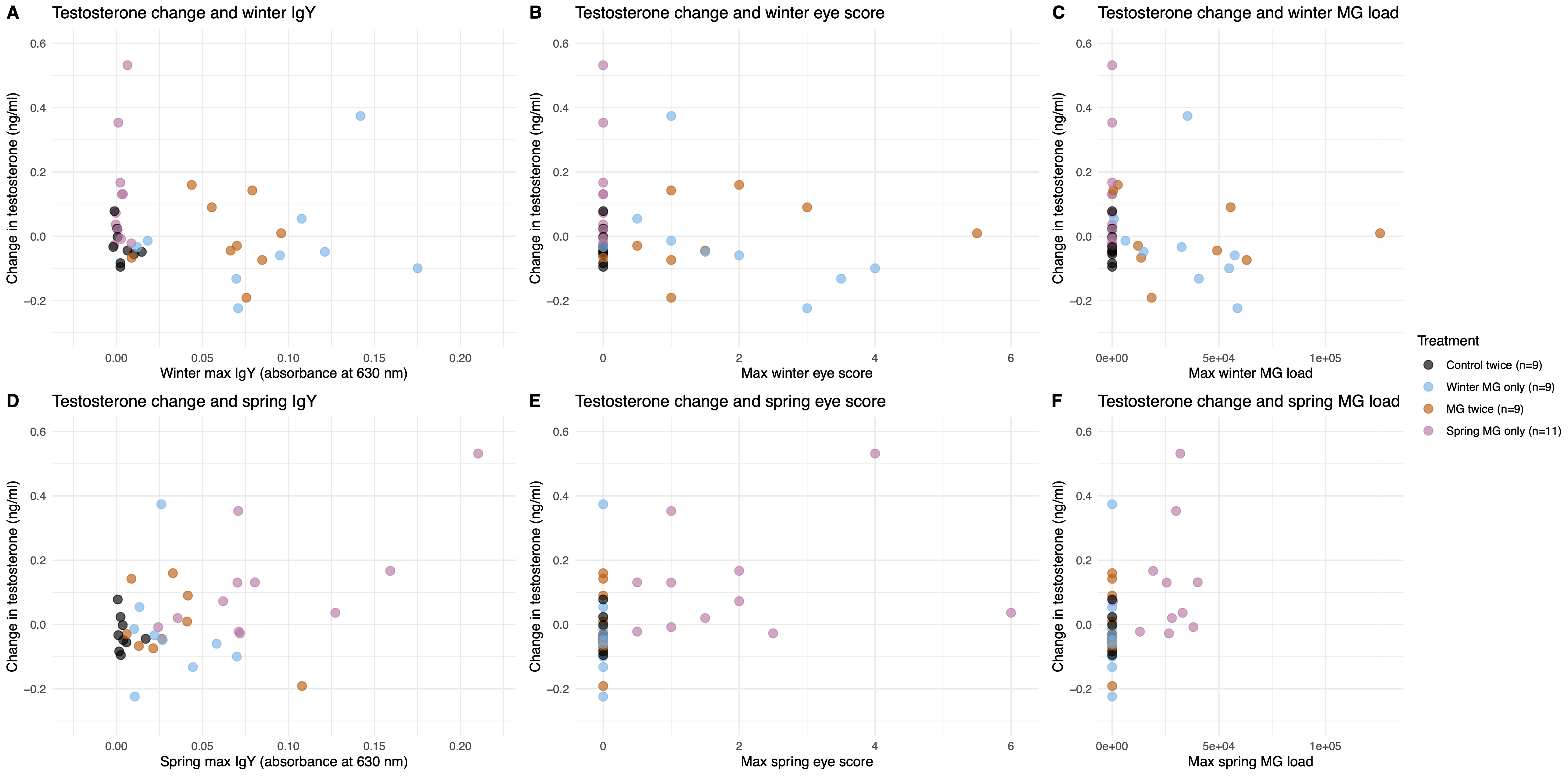

**Figure S6.** Relationships between male house finch (n = 38) testosterone change and responses to *Mycoplasma gallisepticum* (‘MG’), shown by treatment regimen and inoculation response metric. Each bird was first inoculated with either MG or a carrier control while in winter (i.e., non-breeding) condition and then received a second inoculation while in spring (i.e., breeding) condition. Testosterone change was quantified as the increase or decrease in circulating testosterone following spring MG inoculation, compared to winter circulating testosterone; see methods for further quantification details. Shown are relationships between testosterone change and different seasonal metrics of inoculation response. Points are colored by treatment, with finches given two control doses in black, those given MG only in winter in blue, those given two MG doses in orange, and those given MG only during spring in pink.

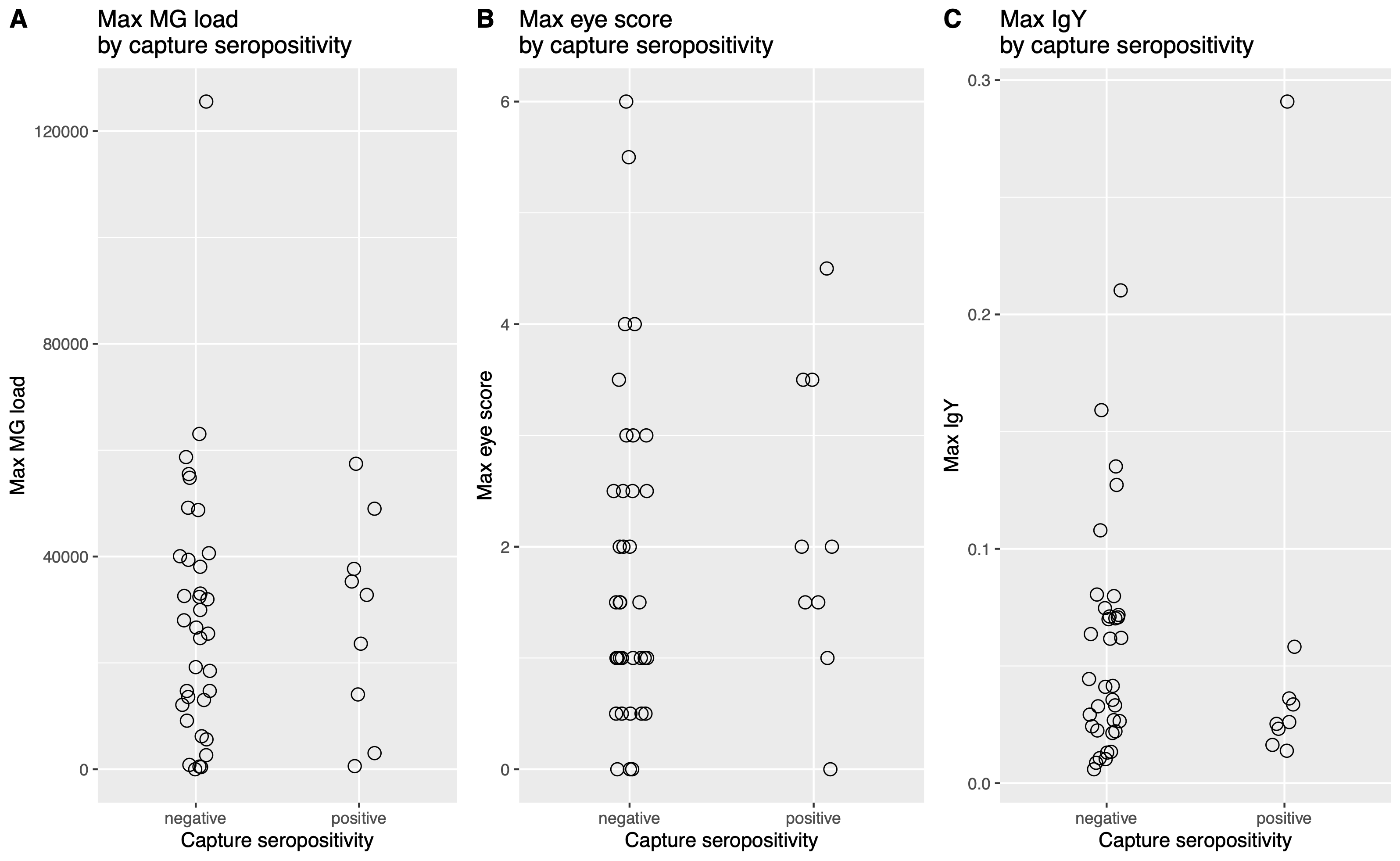

**Figure S7.** Shown are different metrics of inoculation responses (bacterial load, eye pathology score, and antibody production) in house finches inoculated for the first time with *Mycoplasma gallisepticum* (‘MG’), by IgY seropositivity at the point of capture. Points represent data from individual finches that were successfully infected following their first MG inoculation (n = 44) and reflect the maximum observed value for a given metric during the season in which the finch was inoculated for the first time with MG.

**Table S1**. House finches were inoculated with either *Mycoplasma gallisepticum* or a carrier control while in winter (i.e., non-breeding) condition and again while in spring (i.e., breeding) condition. The dimensionality of responses to inoculation (maximum observed anti-MG IgY levels during the winter and spring, maximum observed MG load in winter and spring, and highest eye score observed during winter and spring) was reduced to two dimensions using non-metric multidimensional scaling (NMDS). NMDS loadings are shown for each dimension.

| **Response** | **NMDS1** | **NMDS2** |
| --- | --- | --- |
| Winter IgY | -0.52 | 0.60 |
| Spring IgY | -0.19 | -0.57 |
| Winter MG load | -0.73 | 0.76 |
| Spring MG load | -0.47 | -0.90 |
| Winter eye score | -1.06 | 0.73 |
| Spring eye score | -0.68 | -1.25 |

**Table S2**. Results of PERMANOVA pairwise comparisons of house finch NMDS scores reflecting responses to *Mycoplasma gallisepticum* (‘MG’) treatment. Different treatment regimens are compared by group; experimental treatment (‘Exp’) included inoculation with MG, while control treatment included inoculation with a carrier control (‘Control’). Pairwise differences are based on a Bray-Curtis dissimilarity matrix.

| **Group 1** | | **Group 2** | | **df** | **F** | **R^2^** | **P (Bonferroni)** |
| --- | --- | --- | --- | --- | --- | --- | --- |
| **Winter  treatment** | **Spring treatment** | **Winter  treatment** | **Spring treatment** |  |  |  |  |
| Control | Exp | Control | Control | 1 | 127.07 | 0.82 | 0.01 |
| Control | Exp | Exp | Control | 1 | 107.06 | 0.79 | 0.01 |
| Control | Exp | Exp | Exp | 1 | 68.42 | 0.70 | 0.01 |
| Control | Control | Exp | Control | 1 | 56.17 | 0.68 | 0.01 |
| Control | Control | Exp | Exp | 1 | 52.68 | 0.65 | 0.01 |
| Exp | Control | Exp | Exp | 1 | 1.04 | 0.04 | 1.00 |
