## Supplemental Methods for "Immediate and carryover reproductive costs of infection in female but not male house finches"

#### Finch capture and care

Finches were captured at private residences in and around Blacksburg, VA (Montgomery County, n= 59) and Memphis, TN (Shelby County, n = 1) from May - August 2024. Birds from VA were housed until late September at Virginia Tech in flock cages and provided *ad libitum* water and food (80:20 mixture Roudybush Daily Maintenance Diet, Woodland, CA, USA, and black oil sunflower seeds) and a constant light cycle (12L:12D; hrs light:dark). VA birds were transported to the University of Memphis in an air-conditioned automobile in three IATA-compliant carriers, each with absorbent material, five perches, and ample sunflower seed and water for the journey. No container housed more than 20 birds and all animals were observed for signs of distress (puffing of feathers, panting, lethargy) every four hours during transport. Throughout the experiment, finches were provided with water and food (Daily Maintenance Diet, Roudybush, Woodland, CA, USA) ad libitum. In Memphis, finches were held in flock cages under a 12L:12D photoperiod before being shifted to a winter photoperiod (see methods in main text). All finches were housed singly, with all females housed next to a male neighbor, and remaining males paired with male neighbors. Finches were sexed by plumage coloration at capture and again in Memphis, after their first molt in captivity. Finches with conflicting sex assignments from capture and post-molt were sexed using PCR with primers P2/P8 [1].

*Assessing disease and seropositivity at capture*

Each finch was assessed for clinical signs of MG infection on days 3, 7, and 14 after initial captivity. After 14 days, we drew a small blood sample (<75uL) from the brachial vein to assess levels of MG-specific antibodies using previously published methods [2] and commercially available reagents (FlockChek *M. gallisepticum* ELISA kit, catalog number 99-06729, IDEXX, Westbrook, ME, USA). Briefly, we followed the manufacturer’s instructions, with all steps at room temperature, including an added blocking step: of all wells received 300 μL of 1% bovine serum albumin in phosphate-buffered saline for 40 min. Samples were diluted 1:50 and plated in duplicate; reported values reflect the mean of these duplicates. Absorbance was measured at 630 nm with a microplate reader (ELx800, Bio-Tek, Winooski, VT). MG-specific IgY levels were assessed as sample mean absorbance - dilution-buffer-only mean absorbance. Additional details can be found in [3]. Samples with %CV > 10% were rerun on a new plate. We used 0.061 as our threshold for seropositivity based on the absorbance values for IDEXX assay negative controls. This threshold is conservative, as it is 1.76 SD away from the mean absorbance of negative controls, based on 338 individual wells from 109 plates measured on 79 separate dates across six years.

Most birds (n = 43) showed ELISA results comparable to negative controls (absorbance < 0.06). Birds above this threshold (n = 17) showed absorbance values of between 0.061-0.090 were distributed across treatment groups (7 control winter - control spring; 7 experimental winter - control spring; 2 experimental winter - experimental spring; 1 control winter - experimental spring). Positive control values for these assays ranged from 0.680 - 0.861 (mean = 0.780) and negative control values ranged from 0.041 - 0.044 (mean = 0.042). Capture seropositivity was not associated with load, eye score, or IgY levels following each finch’s first MG inoculation (Figure S7). All finches, except one with an absorbance value of 0.07 (a female that was seronegative at capture), were seronegative by the beginning of the experiment. Because our seropositivity threshold is conservative, it is possible that birds categorized as seropositive at capture had not been previously exposed to MG (i.e., false positives).

#### Photoperiod manipulation

To simulate overwintering, the photoperiod was adjusted to 8L:16D (hrs light:dark) for a period of eight weeks [4] prior to the first MG inoculation. Four days prior to the first inoculation, finches were transferred to paired single-occupancy cages (48.3 x 45.7 x 48.3 cm, each compartment). At 28 days post primary (i.e., winter) inoculation (‘dppi’), finches were placed on long days (14L:10D) to stimulate gonadal recrudescence (modified from [4]).

#### Inoculation protocol and infection success

On winter inoculation day, 18 males and 12 females were inoculated with 20 μL of MG (strain VA94) in Frey’s medium at a concentration of 1.225 x 104 CCU/mL of Frey’s medium, while 20 males and 10 females were inoculated with a carrier control. Doses were administered directly onto the conjunctivae (20 μL per eye) using micropipette droplets. After four weeks of long days, half of each treatment group was inoculated with MG at the same dose as used during winter inoculation, while the other half was inoculated with carrier control; see Figure S1 in the electronic supplementary material for experimental timeline.

Following winter inoculation, all MG-inoculated finches showed evidence of successful infection (i.e., MG load < 50 copies by qPCR, as in [3]) except for three females. No winter controls showed evidence of infection. No finches showed visible signs of infection (i.e., eye score > 0) immediately prior to spring (i.e., second) inoculation. Following spring inoculation, all finches inoculated for the first time were successfully infected, but of the finches inoculated twice, no males were successfully infected, and two of seven females were successfully infected.

#### Sample and data collection

On days -1, 0, 3, 7, 14, 21, 28, 42, and 49 dppi, we collected eye pathology scores (‘eye score’) from each finch. Eye scores are assessed for each eye on an ordinal scale and reflect the degree of conjunctival swelling, from no visible swelling (score of 0) to the eye swollen shut (score of 3); scores for the left and right eye are then summed [5]. On days -1, 3, 7, and 14 dppi, we swabbed the conjunctiva of each eye to quantify MG load [6]. Sterile cotton swabs wetted with tryptose phosphate broth (TPB) were used to swab the interior aspect of both eyes, which was then swirled in a tube of 300 μL TPB to collect conjunctival bacteria. On -1, 14, and 28 days post primary inoculation, we also collected blood samples (< 75 μL) from the wing vein for MG-binding IgY antibodies and hormone quantification. Conjunctival samples and blood samples were held on ice during collection, after which blood samples were centrifuged to separate plasma, and all samples were then stored at -20°C until analysis.

#### MG qPCR

To quantify MG loads from conjunctival samples, we extracted genomic DNA from conjunctival samples by using the Qiagen DNeasy 96 Blood and Tissue kits (Qiagen, Valencia, CA). We used qPCR conditions, primers, probes, and methods outlined in Grodio et al 2008 and updated in Tillman and Adelman 2026. Briefly, each reaction (15 μL total volume) contained 3.525 μL of DNase-free water (Integrated DNA Technologies, Coralville, Iowa, USA), 7.5 μL PrimeTime Gene Expression Master Mix (Integrated DNA Technologies, Coralville, Iowa, USA), 0.375 μL of 10uM forward and reverse primers (Integrated DNA Technologies, Coralville, Iowa, USA), 0.225 μL of 10uM probe (Integrated DNA Technologies, Coralville, Iowa, USA), and 3 μL of template DNA. Assay conditions were: 3 min. at 95℃, then 40 cycles of 3 sec. at 95℃ and 30 sec. at 60℃. We used a Bio-Rad CFX96 (C1000 Touch) thermocycler (Bio-Rad Laboratories, Hercules, CA, USA). Each plate included a serially diluted standard curve (plated in triplicate) of a g-Block (Integrated DNA Technologies, Coralville, IA, USA) based on the mgc2 amplicon from Grodio et al. (2008) ranging from 1.81E+01 to 1.81E+08 copies per well.

#### IgY ELISA

To quantify MG-binding IgY in finch plasma, we used a commercially available enzyme-linked immunosorbent assay (ELISA) kit (FlockChek M. gallisepticum ELISA kit, IDEXX, Westbrook, ME). We quantified anti-MG IgY from blood collected on days -1, 14, and 28 dpsi at University of Memphis using the protocol described above (see ‘Finch capture and care’). IgY levels are reported as sample mean absorbance - dilution-buffer-only mean absorbance, measured at 630 nm.

#### Testosterone ELISA

We quantified circulating testosterone from male plasma samples by using a high sensitivity commercial enzyme immunoassay kit (Enzo Life Sciences ADI-900-176). The precise dilution of each sample depended upon the amount of plasma available, but were, on average 1:17.8 (SD = 6.03, range = 1:81.6 - 1:13.5), with dilutions similar between season (winter: mean = 1:17.1, SD = 7.71, range = 1:81.6 - 1:13.5; spring: mean = 1:18.4, SD = 3.57, range = 1:27.9 - 1:13.5). Only three samples had a dilution of over 1:28. Each diluted sample’s concentration was estimated in pg/mL (see standard curve information below) and was then multiplied by its dilution factor to yield the estimated ng/mL in the original sample. Samples were run in duplicate and the average %CV between replicates (intra-plate variation) was 13.2% for samples above the assay’s limit of detection. Each plate included a standard curve, plated with five standards in duplicate and ranging from 3.9 - 1,000 pg/mL. Plates were read on a BioTek Synergy Plate Reader to assess absorbance at 405nM.

We used a four-parameter-logistic non-linear model in R to generate predicted pg/ML from raw absorbance values of the standards on each plate. Detection limit was determined by assessing where the absorbance value for the lowest replicate of lowest standard intersected with a bootstrapped 95% confidence interval of the standard curve (10,000 iterations) from the *nlstools* package [7] in R. To estimate inter-plate variability, we calculated %CV of the recalculated pg/mL for the second-to-lowest-concentration standard (known concentration = 15.625 pg/mL) across all plates, i.e. that standard’s estimated pg/mL based upon the raw absorbances of its replicate wells and the four-parameter curve described above. This yielded an inter-plate variability of 21.6%. We note, however, that this was heavily influenced by one experiment in which the detection limit was unusually high (23 pg/mL) because of highly variable replicates of the lowest concentration standard. Excluding that experiment, inter-plate variability was 6.8%.

#### Testing model assumptions and fit

For ordinal regressions, we verified the assumption of proportional odds by using a Brant test (*brant* function from the ‘brant’ package [8]. To test ordinal regression fit, we compared each ordinal regression to a null model and generated McFadden’s pseudo R^2^ values using the *pR2* function from the ‘pscl’ package [9]. We confirmed normality of residuals for linear models and generalized linear models (supplemental analyses) by visually inspecting Q-Q plots and using the *DHARMa* package, respectively.
