## Supplemental Analyses for "Immediate and carryover reproductive costs of infection in female but not male house finches"

### Supplementary analyses

*1. Eye score and* Mycoplasma gallisepticum (‘MG’) *load do not vary by sex or season, but finches produce higher IgY when inoculated for the first time during spring*

We used linear models to investigate effects of host sex, season of first inoculation, and the interaction between these, on host responses to initial MG inoculation (peak eye scores and MG load, both square root transformed). Similarly, we used a generalized linear model with a Gamma distribution and log link function to ask whether the same variables predicted IgY levels. All analyses were restricted to winter data for individuals inoculated twice with MG and spring data from individuals inoculated with MG only in the spring; two birds that did not show evidence of successful infection (e.g., eye score > 0 and/or MG load > 15 copies by qPCR) were removed from these analyses (n = 44). Generalized linear model beta estimates were exponentiated to transform them to the original scale of the data.

Among birds inoculated for the first time (n = 44), there was no effect of season, host sex, nor interaction on infection severity as quantified by maximum eye score (F_3,40_ = 0.35, p = 0.79) or maximum MG load (F_3,40_ = 0.56, p = 0.64). However, spring-inoculated finches produced higher levels of IgY than winter-inoculated finches (X^2^ = 30.90, df = 1, p < 0.01, Est(Spring) = 3.25 ± 1.24 SE), although there was no effect of sex. See Figures S2 and S3 in the electronic supplementary information.

#### 2. Higher eye scores are associated with higher bacterial loads, but neither load nor eye score strongly predicts IgY production in Mycoplasma gallisepticum (‘MG’)-infected house finches

Here we use linear models to probe the pairwise relationships among different infection response metrics in house finches inoculated with MG for the first time. Response metrics include maximum observed MG load, maximum observed eye score, and maximum observed IgY. All finches in the dataset were successfully infected (n = 44). Because some finches were inoculated for the first time while in wintering condition (n = 28), and others while in breeding condition (n = 16), we include an interactive effect of season in each model.

Higher maximum MG loads predicted higher eye scores (F = 4.93, df = 3,40, p < 0.01, adjusted R^2^ = 0.22; estimate = 3.36 x 10^-5^ ± 8.86 x 10^-6^ SE, p < 0.01). However, there was no significant interaction between season and maximum MG load on maximum observed eye score (p = 0.36), nor was season a significant predictor of eye score (p = 0.17). There was no significant main effect of MG load (p = 0.58) nor interactive effect of MG and season on IgY (p = 0.54). However, IgY levels were higher in spring than in winter (F = 8.50, df = 3,40, p < 0.01, adjusted R^2^ = 0.34; estimate = 5.70 x 10^-2^ ± 2.85 x 10^-2^ SE, p = 0.05). There was no significant main effect of eye score (p = 0.55) nor interactive effect of season and eye score (p = 0.14) on IgY level. However, there was a trend of higher IgY levels in spring than in winter (F = 10.93, df = 3,40, p > 0.01, adjusted R^2^ = 0.41; estimate = 0.04 ± 0.02 SE, p = 0.10) aligning with the model above. See Figure S3.

#### 3. Spring inoculation responses best predict female reproductive rank

To probe the relationship between female reproductive rank and specific MG inoculation responses, we used ordinal linear regressions, following statistical methods outlined in ‘*Testing model assumptions and fit*’ section of the supplemental methods above. The highest reproductive rank was assigned to females that developed a brood patch and laid eggs, followed by females that developed a brood patch (‘BP’) only, and the lowest rank was given to females showing no signs of reproductive development. Maximum winter IgY (ordinal regression LR = 2.42, df = 1, p = 0.12), eye score (ordinal regression LR = 1.51, df = 1, p = 0.22), and MG load (ordinal regression LR = 0.15, df = 1, p = 0.69) were not significant predictors of reproductive rank; see Figure S5A-C. However, stronger spring inoculation responses were associated with lower odds of laying eggs. Higher spring IgY levels were associated with reduced probability of laying eggs (ordinal regression LR = 24.25, df = 1, p < 0.01, McFadden’s R^2^ = 0.53; estimate = -172.5 ± 66.90), as were higher spring eye scores (ordinal regression LR = 13.10, df = 1, p < 0.01, McFadden’s R^2^ = 0.29; estimate = -36.25 ± 4.18 x 10^-9^) and higher MG loads (ordinal regression LR = 13.83, df = 1, p < 0.01, McFadden’s R^2^ = 0.30; estimate = -0.19 ± 0.13); see Figure S5D-F.

#### 4. Spring inoculation responses best predict male testosterone change

We probed the relationship between testosterone change and specific inoculation responses in male house finches inoculated with *Mycoplasma gallisepticum* (‘MG’). Testosterone change was quantified as the increase or decrease in circulating testosterone following spring MG inoculation, compared to winter circulating testosterone; see methods for further quantification details. Maximum winter IgY (F = 0.51, df = 1,36, p = 0.48), eye score (F = 1.97, df = 1,36, p = 0.17), and MG load (F = 2.19, df = 1,36, p = 0.15) were not predictive of testosterone change; see Figure S6A-C. However, higher maximum spring IgY predicted larger increases in testosterone (F = 10.87, df = 1,36, p < 0.01, adjusted R^2^ = 0.21; estimate = 1.53 ± 0.46 SE, p < 0.01), as did higher maximum spring eye scores (F = 6.91, df = 1,26, p = 0.01, adjusted R^2^ = 0.14; estimate = 0.05 ± 0.02 SE, p = 0.01) and higher maximum spring MG loads (F = 8.97, df = 1,36, p < 0.01, adjusted R^2^ = 0.18; estimate = 4.97 x 10^-6^ ± 1.66 x 10^-6^ SE, p < 0.01); see Figure S6D-F.
